# Co-circulation of Dengue Virus Serotypes 1 and 3 during the 2019 epidemic in Dar es Salaam, Tanzania

**DOI:** 10.1101/763003

**Authors:** Gaspary Mwanyika, Leonard E.G. Mboera, Sima Rugarabamu, Julius Lutwama, Calvin Sindato, Janusz T. Paweska, Gerald Misinzo

## Abstract

**Background:** Dengue is an important mosquito-borne viral disease in tropical and sub-tropical countries. In this study molecular characterization was carried out to determine dengue viruses circulating among patients at health facilities during 2019 epidemic in Dar es Salaam, Tanzania.

**Methods:** The study involved outpatients seeking care for febrile illness at four health facilities in Kinondoni and Ilala Districts of Dar es Salaam City in Tanzania. A total of 45 sera from the outpatients were confirmed dengue-positive for dengue virus (DENV) non-structural protein 1 (NS1) antigen and/or NS1-IgG/IgM antibodies using on-site rapid test. The presence of the virus was detected by reverse-transcription polymerase chain reaction (RT-PCR) method. Of the 45 sera, 20 samples were selected randomly for identification of specific dengue virus serotypes using RT-PCR followed by evaluation of resulting amplicons on agarose gel electrophoresis.

**Findings and significance:** Both Dengue virus serotypes 1 (DENV-1) and 3 (DENV-3) were detected in the samples tested with the former being dominant. We present the first evidence of dengue virus co-infection of DENV-1 and DENV-3 serotypes in Tanzania. The emergence of DENV-1 indicates the possibility of importation of the virus to Tanzania from endemic countries. Due to DENV serotype co-circulation, there is an increased risk of severe dengue in future epidemics. Our findings advocate the importance of genomic-based surveillance to provide rapid evidence of dengue virus emergence/re-emergence and spread.

**Author Summary:** Dengue viruses are the most important mosquito-borne pathogens that pose a serious global health threat. Tanzania has reported several dengue virus epidemics since 2010 with the majority of the epidemics occurring in Dar-es-Salaam city. Until August 2019, a total of 6,859 dengue cases have been confirmed in the country. We performed molecular characterization of dengue viruses (DENV) circulating during the 2019 epidemic phase. It was found that DENV-1 serotype was dominant during the epidemic and two samples of the tested sera were co-infected by DENV-1 and DENV-3 serotypes. These findings emphasize the importance of genomic-based surveillance of dengue viruses in Tanzania to guide strategies for appropriate interventions.

## Introduction

In recent years, Dengue has become the major mosquito-borne viral disease in tropical and subtropical regions affecting people of all age groups in over 128 countries with 50-100 million infections annually [1]. In Africa, dengue is endemic in 34 countries with recent epidemics reported mostly in East Africa [2]. Recent dengue outbreaks have been reported in Tanzania, Mozambique, Ghana, Benin, Ivory Coast and Mauritius [3]. Population growth, increased rate and volume of movement of people, uncontrolled urbanization and climate variability/change have been described to be responsible for global spread of dengue virus (DENV) [4].

In Tanzania, dengue was first described in 1823 by Spanish sailors in the southern coastal region [5]. Since 2010, four dengue outbreaks have been reported in Tanzania [6–9]. To date, the 2019 epidemic seems to be the worst one in the country. From January to August 2019, a total of 6,859 dengue fever cases and 13 deaths have been reported in the country with a large proportion from Dar es Salaam region. Other affected regions include Tanga, Pwani, Lindi, Morogoro, Kagera, Arusha and Ruvuma [9,10].

There are five antigenic distinct serotypes of dengue virus (DENV-1 to 5) [11,12] that can cause mild dengue fever (DF) to life-threatening dengue haemorrhagic fever (DHF) and dengue shock syndrome (DSS) often presented with atypical clinical manifestations like organ impairment [13,14]. Since 1960, four DENV serotypes (DENV-1 to 4) have been reported in Africa with DENV-2 responsible for most of epidemics followed by DENV-1 [15]. In Tanzania, the presence of DENV-3 and DENV-2 serotypes have been documented during the previous epidemics [6–8,15].

Although dengue is endemic in Africa, molecular epidemiology of circulating DENV is not well documented [17,18]. The increasing number of dengue outbreaks and improved access to laboratory diagnostic tools have allowed for more effective recognition of outbreaks through genomic-based surveillance [17,19]. The genomic-based surveillance can help to identify the specific pathogen causing epidemics, track sources and inform the public health authorities for appropriate interventions.

In Tanzania, few studies have reported molecular epidemiology of dengue virus [6,7]. In 2014, Jaswant performed partial sequencing of core pre-membrane junction region of DENV detected in patients during the 2013-2014 outbreak in Dar-es-salaam region and confirmed that DENV-2 serotype genetically close to isolates reported in China and Singapore was responsible for the epidemic [20]. However, genomic-based data are missing for DENV detected in the country during the subsequent epidemics in 2018 and 2019. The objective of this study was to perform molecular characterization of dengue viruses during the 2019 epidemic phase in Dar-es-Salaam, Tanzania.

## Methods

### Study area

This cross-sectional health facility-based study was carried out in Kinondoni and Ilala districts in Dar-es-Salaam, Tanzania (Fig 1). Dar-es-Salaam is usually hot and humid throughout the year with the main dry season from June to September and the short rainy season from October to December followed by long rainy season between March and May. The average daily temperature is 26 °C and total annual rainfall averages 1100 mm with relative humidity of 100% and 60% during the night and day time, respectively [21].

**Fig 1:**
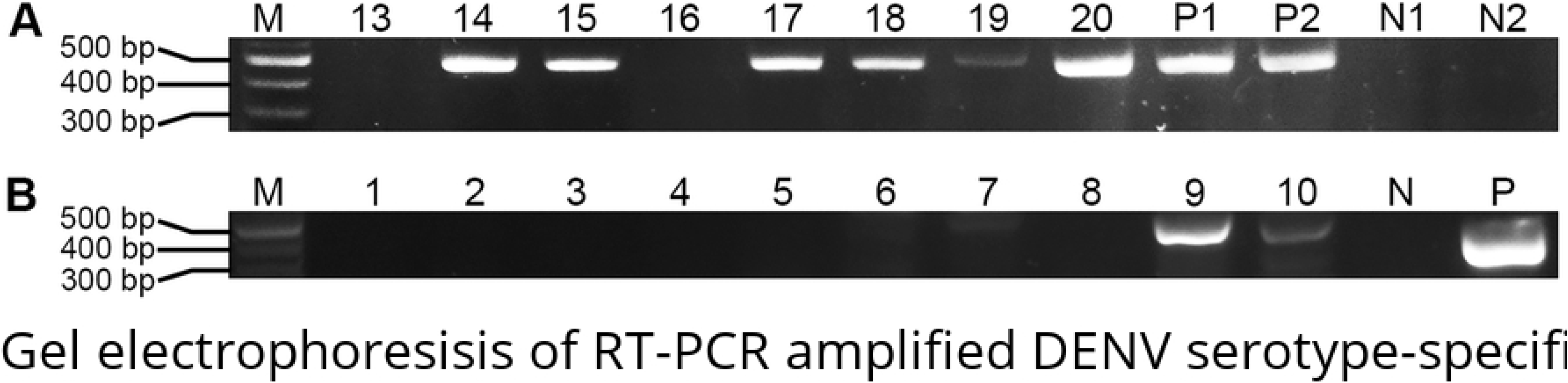
Study location and sample collection sites. Source: This map was designed and drawn in this study

### Patients and sample collection

Dengue-suspected patients were recruited from outpatients at four health facilities, International School of Tanganyika (IST) Clinic and Premier Care Clinic in Kinondoni and Doctors Plaza Hospital and Regency Medical Centre in Ilala district between March and May 2019. A total of 110 dengue-suspected outpatients were enrolled in the study. The inclusion criteria included patients with dengue-like illness and fever (body temperature > 38 °C) prior to onset of clinical manifestations presenting with at least one of the following clinical manifestations namely, retro-orbital pain, rash, arthralgia, malaise, signs of haemorrhaging and organ failure. Fever patients with bacterial infections and those unwilling to participate in this study were excluded. Haematology data were obtained from routine blood tests performed by the hospitals. All dengue positive cases were categorized clinically either as dengue fever (DF), dengue haemorrhagic fever (DHF), dengue shock syndrome (DSS), or expanded dengue syndrome (EDS), according to the World Health Organization guidelines [22].

Peripheral venous blood was collected into serum separator tubes for serum separation and testing. The on-site duo dengue NS1-IgG/IgM rapid test (CTK BIOTECH Inc, CA, USA) was used to detect the presence of NS1 antigen and IgG/IgM antibodies against DENV NS1. Aliquots of a total of 45 positive sera samples were kept in well-labeled cryotubes stored at −80 °C. Thereafter, sera samples were transported in a cool box with dry-ice to Sokoine University of Agriculture (SUA) molecular biology laboratory in Morogoro and kept at −80 °C until genomic analysis.

### RNA extraction

Out of 45 sera samples, we focused on 20 samples that were selected randomly. A total of 60 μL viral nucleic acid (RNA) was extracted from 140 μL of each test sample using QIAamp RNA mini kit (Qiagen, Hilden, Germany) following the manufacturer’s instructions. The extracted RNA was stored immediately in aliquots at −20 °C to avoid freeze-thawing cycles that could damage RNA stability. We also assessed the quality of extracted RNA by measuring absorbance at 260 and 280 nm using the NanoDrop ND1000 spectrophotometer (GE Healthcare UK Limited, Buckinghamshire, UK). The A260/280 ratio of ~2.0 was accepted as pure RNA [23].

### Detection of DENV RNA

Dengue virus nucleic acid (RNA) was detected by RT-PCR using SuperScript III Platinum *Taq* DNA polymerase (Invitrogen, Carlsbad, CA, USA) One-Step RT-PCR following the manufacturer’s instructions. RT-PCR amplification was performed with primers as previously described [24]. We prepared a 25 μL RT-PCR reaction containing 12.5 μL of 2x reaction mix, 1 μL of SuperScript III RT/Platinum *Taq* mix, 0.5 μL of 10 μM sense primer (D1), 0.5 μL of 10 μM anti-sense primer (D2), 0.5 μL Magnesium salt (Invitrogen, CA, USA), 6 μL of nuclease-free water and 4 μL of RNA template. Reverse-transcription reaction was performed in 1 cycle at 48 °C for 30 minutes, followed by one cycle of initial denaturation at 94 °C for 2 minutes. PCR amplification was done for 35 cycles of denaturation at 94 °C for 15 seconds, annealing at 55 °C for 30 seconds and elongation at 68 °C for 1 minute. This was followed by a final extension at 68 °C for 5 minutes. The RT-PCR products with the expected band size 511 base pairs (bp) were evaluated on 1.5% agarose gel stained with Gel Red (Biotium Inc, CA, USA) against 100 bp DNA ladder (Bio-Rad Laboratories, Hercules, CA, USA) at 100 V for 45 minutes. The specific PCR bands were visualized using Gel Doc EZ Imager system (Bio-Rad Laboratories, Hercules, CA, USA).

### Identification of DENV serotypes

Reverse-transcription polymerase chain reaction (RT-PCR) was performed to detect DENV serotypes using serotype-specific primers. The amplification of serotype-specific nucleotide sequences was done in a 25 μL RT-PCR reaction of each serotype test containing 12.5 μL of 2x reaction mix, 1 μL of SuperScript III RT/Platinum *Taq* mix (Invitrogen, CA, USA), 0.5 μL of 10 μM forward primer (D1), 0.5 μL of 10 μM reverse primer (TS1/TS2/TS3/TS4) (Table 1), 0.5 μL Magnesium salt (Invitrogen, CA, USA), 6 μL of nuclease-free water and 4 μL of RNA template. Reverse-transcription was performed in 1 cycle at 50 °C for 10 minutes, followed by initial denaturation at 94 °C for 2 minutes. PCR was done for 35 cycles of denaturation at 94 °C for 15 seconds, annealing at 55 °C for 30 seconds, elongation at 72 °C 1 minute and a final extension at 72 °C for 7 minutes. The expected sizes of serotype-specific amplicons were 482 bp (DENV-1), 119 bp (DENV-2), 290 bp (DENV-3) and 392 bp (DENV-4). The PCR bands were analysed on 1.5% agarose gel against 100 bp DNA ladder (Bio-Rad Laboratories, Hercules, CA, USA) at 100 V for 45 min and visualized using Gel Doc EZ Imager system (Bio-Rad Laboratories, Hercules, CA, USA).

**Table 1:**
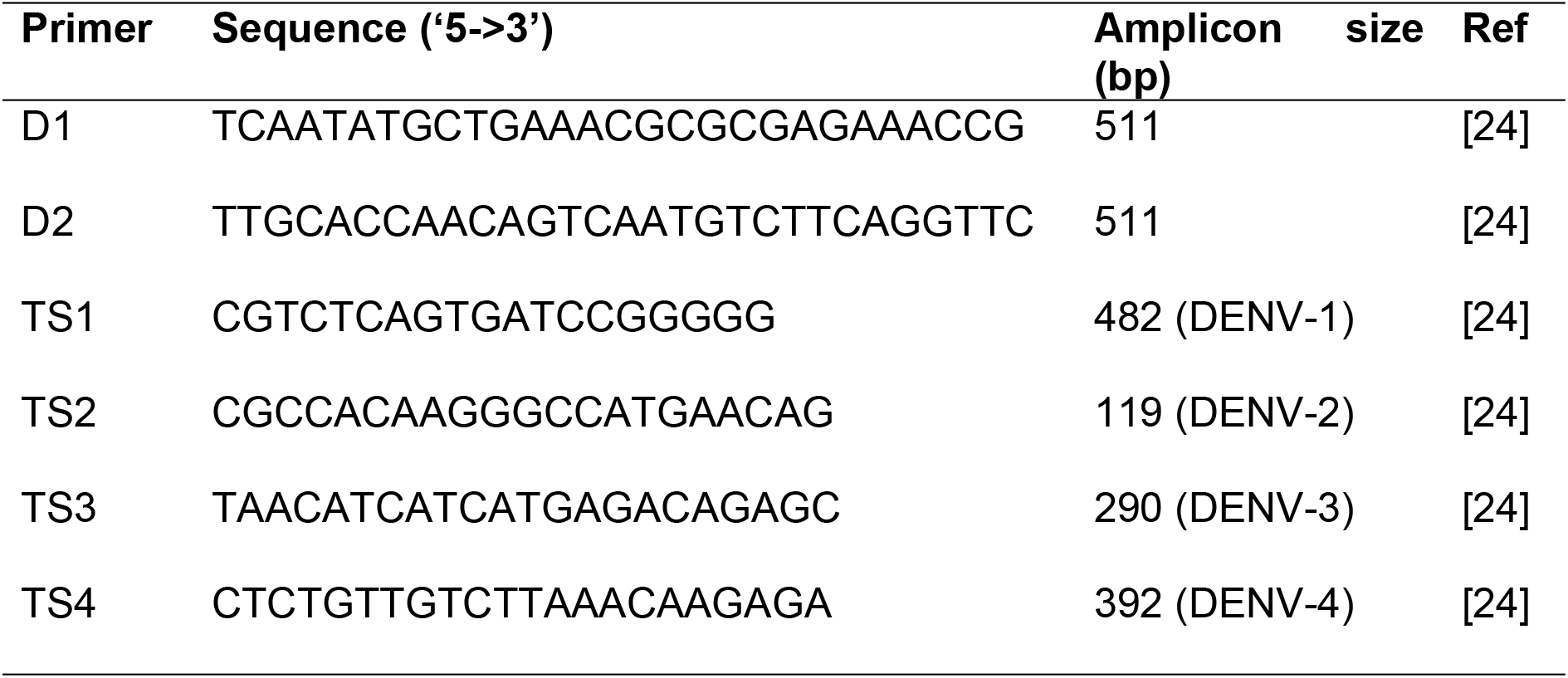
Primers used in RT-PCR amplification.

### Data analysis

Socio-demographic data were entered in Miscrosoft Execel spreadsheet (MS-Excel 2010, Microsoft Corp., and Redmond, WA, USA) and descriptive statistics [counts, percentages, median and interquartile range (IQR)] and relevant tables were used to summarize information. RT-PCR bands were analysed using Image Lab Software version 6.0.1 (Bio-Rad Laboratories, Hercules, CA, USA) and presented by gel images.

### Ethical consideration

The study was approved by the Medical Research Coordinating Committee of the National Institute for Medical Research in Tanzania (Ref. No. NIMR/HQ/R.8a/Vol.IX/2974). Witten consent was sought and obtained from all adults (≥ 18 years) and assent was obtained from parents/guardians of children (<18years old) for screening of dengue virus infection. Oral consent was sought from dengue-positive individuals for further identification and genomic analysis and it was documented through tracking forms.

## Results

### Patient characteristics and dengue diagnosis

A total of 110 sera samples was collected from dengue-suspected outpatients presented at health facilities with dengue-like illiness. Among them, 45 (41%) were confirmed positive for dengue infection by NS1 antigen and/or DENV RNA detection with RT-PCR (95.5% positivity by RT-PCR and/or NS1 and 4.5% positivity by NS1 only). The age of the patients ranged from 16-75 years old (median = 35, IQR= 18). Two-thirds of the patients (66.7%) were males; 62.2% had post-secondary education and the majority (46.7%) had formal employment (Table 2).

**Table 2:** Socio-demographic characteristics of the confirmed dengue-positive patients (n=45)

| Characteristic | Total participants (%) |
| --- | --- |
| Age group |  |
| < 18 | 4 (8.9) |
| 18-24 | 5 (11.1) |
| 25-34 | 9 (20) |
| 35-44 | 14 (31.1) |
| 45-54 | 6 (13.3) |
| 55 and above | 7(15.6) |
| Sex |  |
| Male | 30 (66.7) |
| Female | 15 (33.3) |
| Education |  |
| None | 2 (4.4) |
| Primary | 1 (2.2) |
| Secondary | 14 (31.1) |
| College/University | 28 (62.2) |
| Occupation |  |
| Self employed | 13 (28.9) |
| Business | 4 (8.8) |
| Formal employment | 21 (46.7) |
| Students | 7(15.6) |

### Detection of DENV RNA and serotypes

Dengue virus (DENV) nucleic acid (RNA) was detected in 17 of 20 tested samples while three samples were negative. Serotype-specific RT-PCR test was positive for DENV-1 (Fig 2A) and DENV-3 (Fig 2B) serotypes. Of the 17 positive samples, 17 were positive for DENV-1, of which two samples were also positive for DENV-3. RT-PCR results for DENV-2 and DENV-4 serotypes were all negative. The overall results for RNA quality and RT-PCR amplification of the tested samples are summarized in Table 3.

**Fig 2:**
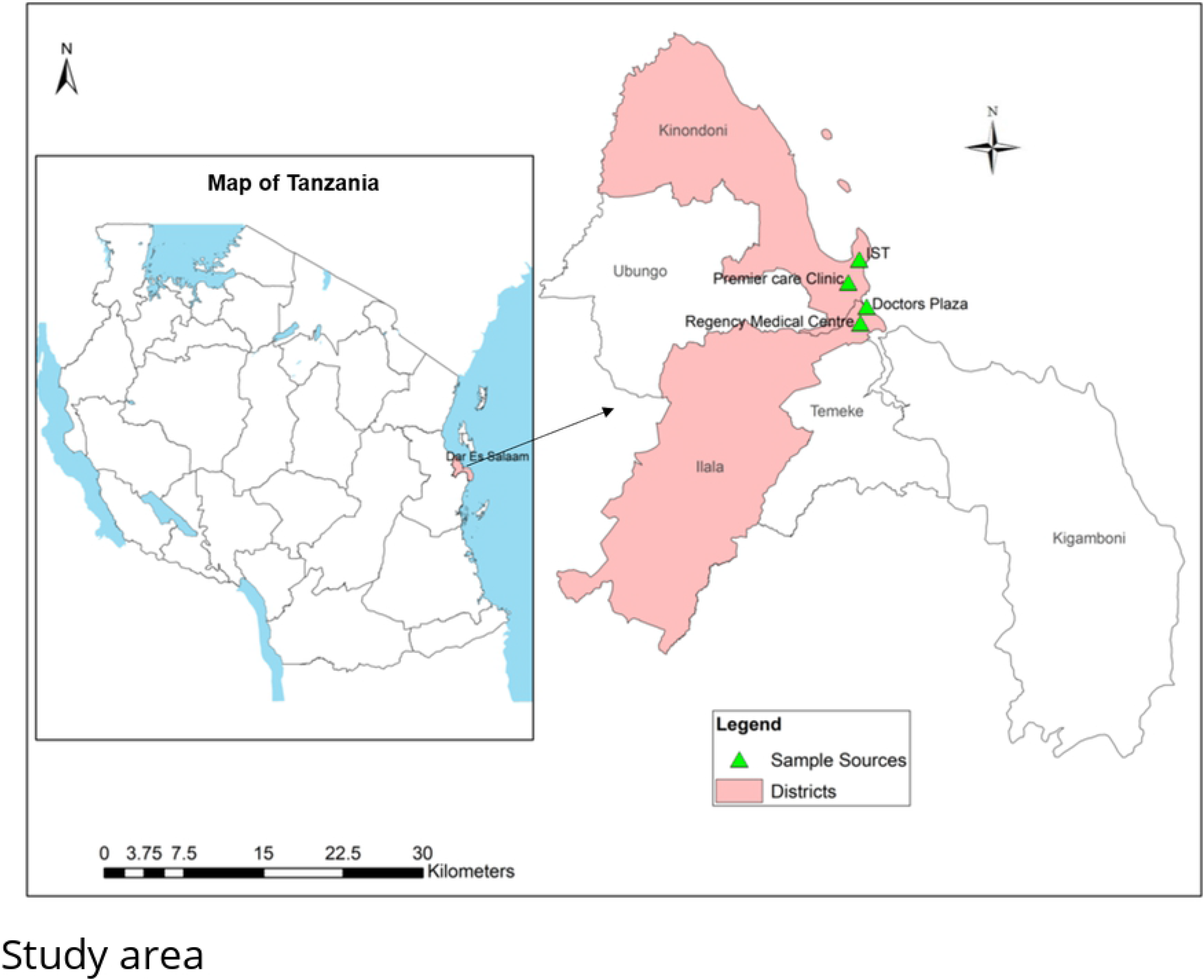
Agarose gel electrophoresis image for RT-PCR amplified serotype-specific DENV DNA products. (A) Dengue virus serotype 1 (482 bp), M is a 100 bp DNA ladder, 13-20 are test sera samples, P1, P2 are positive controls (CDC KK0129) and N1, N2 are the negative controls. (B) Dengue virus serotype 3 (290 bp), M is a 100 bp DNA ladder, 1-10 are test sera samples, N is the negative control and P is positive control (CDC KK0129).

**Table 3:** RNA quality and RT-PCR results of the tested sera (n=20)

| Sample ID | District | *Hospital | Sex | Age | RNA quality (A260/280) | DENV RNA | Serotype |
| --- | --- | --- | --- | --- | --- | --- | --- |
| 1 | Kinondoni | RMC | M | 16 | 2.83 | +<sup>a</sup> | DENV-1 |
| 2 | Kinondoni | RMC | M | 48 | 2.25 | + | DENV-1 |
| 3 | Ilala | DPH | M | 45 | 1.54 | + | DENV-1 |
| 4 | Kinondoni | IST | F | 34 | 2.75 | + | DENV-1 |
| 5 | Kinondoni | DPH | F | 36 | 1.42 | -<sup>b</sup> | N/A<sup>c</sup> |
| 6 | Temeke | RMC | M | 56 | 2.09 | + | DENV-1 |
| 7 | Temeke | RMC | M | 26 | 2.45 | + | DENV-1 |
| 8 | Kinondoni | DPH | M | 35 | 2.22 | + | DENV-1 |
| 9 | Kinondoni | DPHC | M | 44 | 1.57 | + | DENV-1/3<sup>d</sup> |
| 10 | Temeke | PC | M | 40 | 1.66 | + | DENV-1/3 |
| 11 | Ilala | PC | M | 65 | 4.98 | + | DENV-1 |
| 12 | Kinondoni | IST | M | 29 | 5.39 | + | DENV-1 |
| 13 | Ilala | RMC | M | 22 | 1.69 | - | N/A |
| 14 | Kinondoni | DPH | M | 33 | 2.81 | + | DENV-1 |
| 15 | Kinondoni | DPH | M | 20 | 2.61 | + | DENV-1 |
| 16 | Kinondoni | PC | M | 27 | 1.53 | - | N/A |
| 17 | Kinondoni | PC | M | 21 | 1.95 | + | DENV-1 |
| 18 | Ilala | IST | F | 37 | 2.00 | + | DENV-1 |
| 19 | Kinondoni | RMC | M | 21 | 2.14 | + | DENV-1 |
| 20 | Kinondoni | RMC | F | 31 | 2.43 | + | DENV-1 |

<sup>a</sup>Positive for dengue virus nucleic acid (DENV RNA) reverse-transcription polymerase chain reaction test (RT-PCR); <sup>b</sup> Negative for DENV RNA-RT-PCR test; <sup>c</sup>N/A, Not applicable; <sup>d</sup>Positive for DENV serotypes 1 and 3 RT-PCR test; \*RMC, Regency Medical Centre; DPH, Doctors Plaza Hospital; IST, International School of Tanganyika Clinic and PC, Premier Care Clinic.

## Discussion

This study reports for the first time occurrence of dengue virus serotype 1 (DENV-1) in Tanzania. The occurrence of DENV-2 and DENV-3 has been reported during the previous outbreaks in 2010 and 2014 [6,7,25]. Our results indicate that DENV-1 was dominant serotype during the 2019 outbreak in Tanzania. DENV-1 has also been detected in Japan in a patient with a history of travel to Tanzania in May 2019 [26]. The subsequent phylogenetic analysis indicated that the DENV-1 isolate detected in the Japanese traveller was genetically related to Singapore 2015 strain, suggesting that the recent DENV-1 epidemic in Tanzania may have been imported from endemic countries.

The evidence of DENV-1 predominance in the East Africa region is increasingly been reported. In a recent study in Kenya, DENV-1 was found to account for 44% of all the serotypes that were detected in febrile patients [27]. The results from this study demonstrated that DENV-1 isolates were genetically close to isolates reported in endemic South-East Asian countries like Indonesia and India. Nevertheless, the risk of DENV-1 serotype importation to Tanzania can originate from countries other than South-East Asian countries due to increased movement networks of people. For instance, Mauritius has recently reported DENV-1 epidemic that included eleven imported cases from South-East Asia countries [9]. Circulation of DENV-1 serotypes in Africa has been previously reported in Cameroon, Comoros, Côte d’Ivoire, Djibouti, Gabon, Kenya, and Senegal [2,28].

Also, we report for the first time genomic-based evidence of dengue virus co-infection with DENV-1 and DENV-3 serotypes during a single phase of outbreak in Tanzania. Dengue concurrent infections have been commonly reported in endemic countries including Brazil [29] and India [30]. Dengue concurrent infections have serious clinical implication for patient management, as they can present with severe clinical manifestations such as haemorrhagic fever syndrome [31,32]. Previous findings showed that subsequent infection with a different serotype may lead to increased disease severity because antibodies against the previous serotype do not cross-neutralize the subsequent heterologous invading serotype, but induce antibody-dependent enhancement that increases replication inside target Fc receptor-bearing cells [33,34]. Co-infection with multiple DENV serotypes may increase the risk of emergence of recombinant strains that have different characteristics and the areas with a circulation of more than one serotype have been classified as more prone to severe DENV infections [31,35]. Co-circulation of dengue virus serotypes and occurrence of serotypes co-infection in Tanzania provide alert on the possibility of an increased risk of severe dengue in the country.

It is worth noting important limitations of this study. This was a cross-sectional health facility-based study that focused on the detection of dengue viruses from outpatients during an epidemic phase, thus, it was not possible to establish an algorithm to classify primary and secondary dengue virus infections and sequence DENV RNA genomes that could provide insights on the evolution of circulating dengue viruses. Also, the health facility-based subjects included in the study may not be a representative of the population parameter estimates, suggesting that extrapolation of the results to unsampled locations should be made with caution.

In conclusion, the findings of this study indicate that DENV-1 is the dominant serotype during the on-going 2019 epidemic and reveal for the first time occurrence of dengue virus serotypes 1 and 3 co-concurrent infection in Tanzania. These observations emphasize the need for establishing continuous genomic-based surveillance of circulating dengue viruses in Tanzania. This will provide rapid evidence of emerging/re-emerging and spread of dengue viruses and inform public health authorities to implement effective preventive measures.

## Acknowledgments

We are grateful to the Management of Regency Medical Centre, Doctor’s Plaza Hospital, International School of Tanganyika Clinic and Premier Care Clinic for providing sera samples used in this study. Lancet Laboratories Tanzania is thanked for laboratory technical assistance in serum separation and testing. We would like to thank Ms Mariam Makange of the Sokoine University of Agriculture Molecular Laboratory for her technical support during molecular laboratory analysis. We also, acknowledge the support from the Government of Tanzania and the World Back through the SACIDS-Africa Centre of Excellence for Infectious Disease of Human and Animal.

